# EMPress enables tree-guided, interactive, and exploratory analyses of multi-omic datasets

**DOI:** 10.1101/2020.10.06.327080

**Authors:** Kalen Cantrell, Marcus W. Fedarko, Gibraan Rahman, Daniel McDonald, Yimeng Yang, Thant Zaw, Antonio Gonzalez, Stefan Janssen, Mehrbod Estaki, Niina Haiminen, Kristen L. Beck, Qiyun Zhu, Erfan Sayyari, Jamie Morton, Anupriya Tripathi, Julia M. Gauglitz, Clarisse Marotz, Nathaniel L. Matteson, Cameron Martino, Jon G. Sanders, Anna Paola Carrieri, Se Jin Song, Austin D. Swafford, Pieter C. Dorrestein, Kristian G. Andersen, Laxmi Parida, Ho-Cheol Kim, Yoshiki Vázquez-Baeza, Rob Knight

## Abstract

Standard workflows for analyzing microbiomes often include the creation and curation of phylogenetic trees. Here we present EMPress, an interactive tool for visualizing trees in the context of microbiome, metabolome, etc. community data scalable beyond modern large datasets like the Earth Microbiome Project. EMPress provides novel functionality—including ordination integration and animations—alongside many standard tree visualization features, and thus simplifies exploratory analyses of many forms of ‘omic data.

## Main Text

The increased availability of sequencing technologies and automation of molecular methods have enabled studies of unprecedented scale [1] prompting the creation of tools better suited to store, analyze [2], and visualize [3] studies of this magnitude. Many of these tools, such as [4, 5, 6, 7], use phylogenies detailing the evolutionary relationships among features or dendrograms that organize features in a hierarchical structure (e.g. clustering of mass spectra) [8]. The challenge of enabling fully interactive analyses stems from the disconnect between feature-level tools and dataset-level tools; few can interactively integrate multiple representations of the data [9], and to our knowledge none scale to display large datasets. This is a key unresolved challenge for the field: to allow researchers to contextualize community-level patterns (groupings of samples) together with feature-level structure, i.e. which features lead to the groupings explained in a given sample set.

Here, we introduce EMPress (https://github.com/biocore/empress), an open-source (BSD 3-clause), interactive and scalable phylogenetic tree viewer accessible as a QIIME 2 [2] plugin. EMPress is built around the high-performance balanced parentheses tree data structure [10], and uses a hardware-accelerated WebGL-based rendering engine that allows EMPress to visualize trees with hundreds of thousands of nodes using a laptop’s web browser (Methods). By integrating EMPress with the widely-used EMPeror software [3] within QIIME 2, EMPress can simultaneously visualize a phylogenetic tree of features in a study coupled with an ordination of the same study’s samples. User actions in one visualization, such as selecting a set of samples in the ordination, update the other, providing context that would not be easily accessible with independent visualizations. This tight integration between displays streamlines several use-cases elaborated below that previously required manual investigation or writing custom scripts.

EMPress visualizations can be created solely from a tree, or users can provide additional metadata files and a feature table to augment the tree. Using these common data files,users can interactively configure many visual attributes in the tree (see Methods and Figures for examples).

Rather than providing a programmatic interface for the procedural generation of styled phylogenetic trees [11, 12, FigTree (http://tree.bio.ed.ac.uk/software/figtree/)], EMPress provides an interactive environment to support exploratory feature- and sample-level tree-based analyses. Many use-cases supported in EMPress accommodate community analysis tasks; this differs from Anvi’o [13] which is centered on the analysis of metagenomic assembled-genomes, pangenomes, etc.. PHYLOViZ [9], SigTree [14], and iTOL [15] are similar to EMPress in terms of their implementation (PHYLOViZ Online also uses WebGL), and/or use-cases (SigTree is mostly used to visualize differential abundance patterns, and iTOL supports the visualization of QIIME 2 tree artifacts). EMPress stands out in its scalability: iTOL claims trees with more than 10,000 tips to be “very large” (https://itol.embl.de/help.cgi), while EMPress readily supports trees with over hundreds of thousands of tips, as shown in Fig. 1. Many visualization customization options available in EMPeror, iTOL [15] and Anvi’o [13] are immediately accessible in EMPress’ interface. Continuous feature metadata can be visualized in tip-level barplots as a color gradient and/or by adjusting the lengths of individual tips’ barplots; categorical sample metadata information can be visualized using a stacked barplot showing—for each tip—the proportion of samples containing that tip stratified by category. These options are available on the user interface and do not require programming or configuration files.

**Figure 1.**
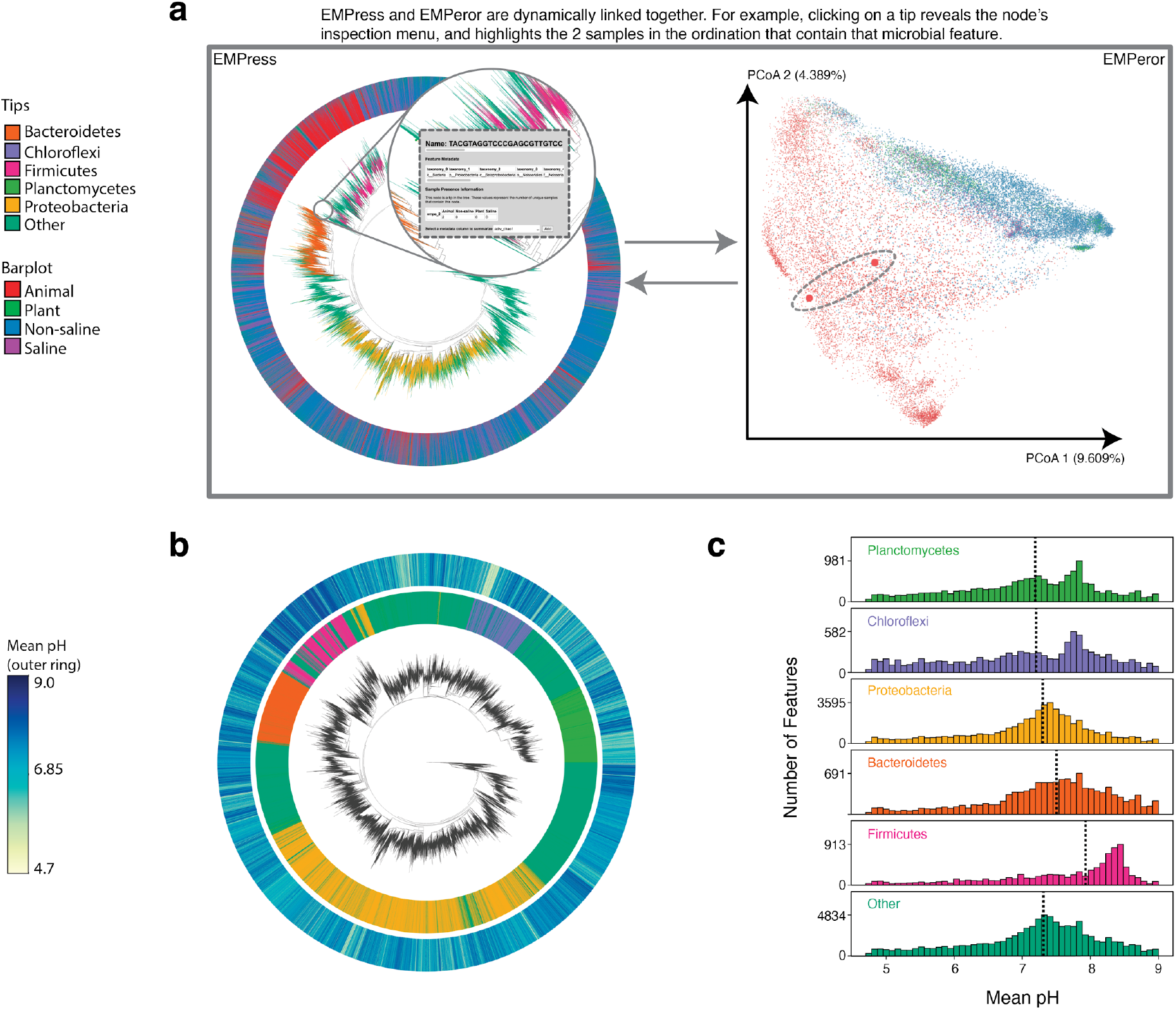
Earth Microbiome Project paired phylogenetic tree (including 756,377 nodes) and unweighted UniFrac ordination (including 26,035 samples). **(a)** Graphical depiction of Empress’ unified interface with fragment insertion tree (left), and unweighted UniFrac sample ordination (right). Tips are colored by their phylum-level taxonomic assignment; the barplot layer is a stacked barplot describing the proportions of samples containing each tip summarized by level 2 of the EMP ontology. Inset shows summarized sample information for a selected feature. The ordination highlights the two samples containing the tip selected in the tree enlarged to show their location. **(b)** Subset of EMP samples with pH information: the inner barplot ring shows the phylum-level taxonomic assignment, and the outer barplot ring represents the mean pH of all the samples where each tip was observed **(c)** pH distributions summarized by phylum-level assignment with median pH indicated by dotted lines. Interactive figures can be accessed <u>here</u>.

Ordination plots computed from UniFrac distances are often used to visualize sample clustering patterns in microbiome studies. However, interpreting the patterns in these plots—and determining which features influence sample group separation—is not always straightforward. While biplots show information about influential features alongside samples, the phylogenetic relationships of these features are not immediately obvious. EMPress aids interpretation of these plots by optionally providing a unified interface where the tree and ordination visualizations are displayed side-by-side and “linked” through sample and feature identifiers [16]. This combination allows for novel exploratory data analysis tasks. For example, selecting a group of samples in the ordination highlights nodes in the tree present in those samples, and vice versa (see Methods). This integration extends to biplots: clicking feature arrows in the ordination highlights their placement in the tree. Lastly, EMPress allows visualizing longitudinal studies by simultaneously showing the tree nodes unique to groups of samples at each individual time point during an EMPeror animation (see Methods).

Using the first data release of the Earth Microbiome Project (EMP), we demonstrate EMPress’ scalability by rendering a 26,035 sample ordination and a 756,377 node tree (Figure 1A). To visualize the relative proportions of taxonomic groups at the phylum level, we use EMPress’ feature metadata coloring to highlight the top 5 most prevalent phyla (see Methods). Next, we add a barplot layer showing, for each tip in the tree, the proportions of samples containing each tip summarized by level 2 of the EMP ontology (Animal, Plant, Non-Saline, and Saline). Paired visualizations allow us to click on a tip in the tree and view the samples that contain that feature in the ordination. This functionality is useful when analyzing datasets with outliers or mislabeled metadata. Tip-aligned barplots summarize environmental metadata: for example, Figure 1B shows the subset of samples (4,002) with recorded pH information and a barplot layer with the mean pH where each feature was found. The barplot reveals a relatively dark section near many Firmicutes-classified features on the tree; in concert with histograms showing mean pH for each phylum (Figure 1C), we can confirm that Firmicutes-classified features are more commonly found in higher pH environments.

EMPress can be applied to various ‘omic datasets. To illustrate this versatility we reanalyzed a COVID-19 metatranscriptome sequencing dataset [17], a liquid chromatography mass-spectrometry (LC-MS) untargeted metabolomic food-associated dataset [8], and a 16S rRNA sequencing oral microbiome dataset [18]. Despite the vastly different natures of these datasets, EMPress provides meaningful functionality for their analysis and visualization. Supplemental Video 1 (<u>supplementary-video-1.mp4</u>) shows a longitudinal exploratory analysis using EMPress and EMPeror representing a subset of SARS-CoV-2 genome data from GISAID. This paired visualization emphasizes the relationships in time and space among “community samples” and the convergence of locales in the United States with the outbreak in Italy (See Methods). The interactive nature of EMPress allows rapid visualization of strains observed in a collection of samples from different geographical locations.

Figure 2A showcases Empress’ ability to identify feature clusters that are differentially abundant in COVID-19 patients compared to community-acquired pneumonia patients and healthy controls [17]. Clades showing KEGG enzyme code (EC) [19] annotations are collapsed at level two except for lyases, highlighting feature 4.1.1.20 (carboxy-lyase diaminopimelate decarboxylase) that was more abundant in COVID-19 here and in an independent metaproteomic analysis of COVID-19 respiratory microbiomes [20].

**Figure 2.**
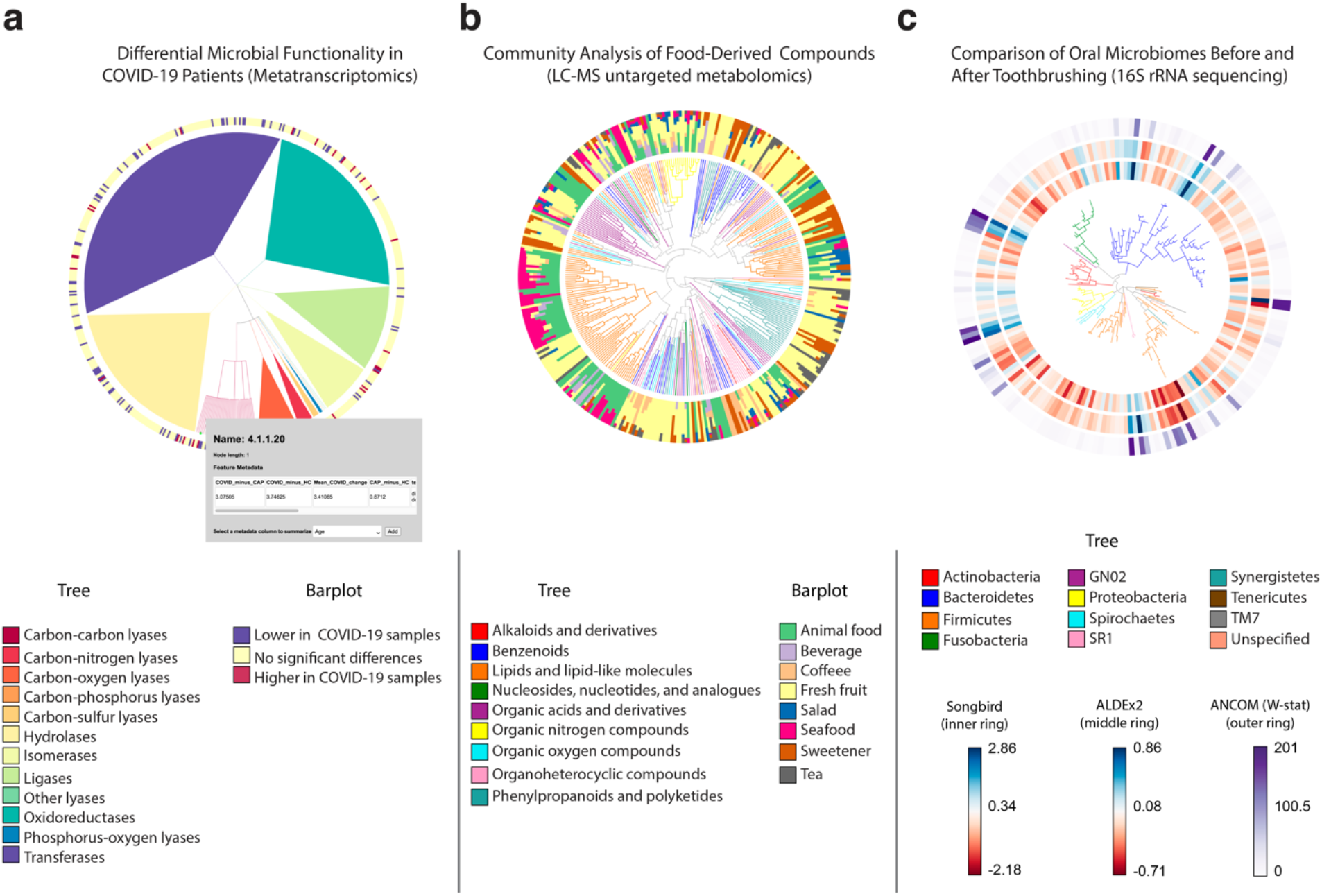
EMPress is a versatile exploratory analysis tool adaptable to various -omics data types. **(a)** RoDEO differential abundance scores of microbial functions from metatranscriptomic sequencing of COVID-19 patients (n=8), community-acquired pneumonia patients (n=25), and healthy control subjects (n=20). The tree represents the four-level hierarchy of the KEGG enzyme code. The barplot colors significantly differentially abundant features (p<0.05) in COVID-19 patients. Clicking on a tip produces a pop-up insert tabulating the name of the feature, its hierarchical ranks, and any feature annotations. **(b)**Global FoodOmics Project LC-MS data. Stacked barplots indicate the proportions of samples (n=70) (stratified by food) containing the tips in an LC-MS Qemistree of food-associated compounds, with tip nodes colored by their chemical superclass. **(c)** *de novo* tree constructed from 16S rRNA sequencing data from 32 oral microbiome samples. Samples were taken before (n=16) and after (n=16) subjects (n=10) brushed their teeth; each barplot layer represents a different differential abundance method’s measure of change between before- and after-brushing samples. The innermost layer shows estimated log-fold changes produced by Songbird; the middle layer shows effect sizes produced by ALDEx2; and the outermost layer shows the W-statistic values produced by ANCOM (see Methods). The tree is colored by tip nodes’ phylum-level taxonomic classifications. Interactive figures can be accessed <u>here</u>.

Recent developments in cheminformatics enabled the analysis and visualization of small molecules in the context of a cladogram [8]. Using a tree that links molecules by their structural relatedness, we analyzed untargeted LC-MS/MS data from 70 food samples (see Methods). With EMPress’ sample metadata barplots, we can inspect the relationship between chemical annotations and food types. Figure 2B shows a tree where each tip is colored by its chemical super class, and where barplots show the proportion of samples in the study containing each compound by food type. This representation reveals a clade of lipids and lipid-like molecules that are well represented in animal food types and seafoods. In contrast, salads and fruits are broadly spread throughout the cladogram.

Lastly, in Figure 2C, we compare three differential abundance methods in an oral microbiome dataset [18] as separate barplot layers on a tree. This dataset includes samples (n=32) taken before and after subjects brushed their teeth (see Methods). As observed across the three differential abundance tools’ outputs, all methods agree broadly on which features are particularly “differential” (for example, the cluster of Firmicutes-classified sequences in the bottom-right of the tree; see Methods), although there are discrepancies due to different methods’ assumptions and biases.

## Conclusions

By providing an intuitive interface supporting both categorically new and established functionality, EMPress complements and extends the available range of tree visualization software. EMPress can perform community analyses across distinct “omics” types, as demonstrated here. Moving forward, facilitating the integration of multiple orthogonal views of a dataset at a more generalized framework level (for example, using QIIME2’s [2] visualization API) will be important as datasets continue to grow in complexity, size, and heterogeneity.

## Acknowledgements

We thank members of the Knight Lab and IBM AIHL Bioinformatics team for feedback during code reviews and presentations. We gratefully acknowledge the following Authors from the originating laboratories responsible for obtaining the specimens and the submitting laboratories where genetic sequence data were generated and shared via the GISAID Initiative, on which a portion of this research is based (Supplemental Table 1).

## Funding

This work is partly supported by IBM Research AI through the AI Horizons Network and the UC San Diego Center for Microbiome Innovation (to KC, MWF, JM, QZ, TZ, JS, ADS, SS, YVB, RK); CCF foundation #675191 (RK and PCD), U19 AG063744 01 (RK, JMG, PCD), CDC contract #75D30120C09795 (KGA and RK), U19 AI135995 (KGA).

## Author Contributions

KC, QZ, YY, JM, TZ, JS, and RK conceived the original idea for the project. KC, MWF, GR, DM, AG, SJ, ME, YY, ES, JM, TZ, QZ, YVB wrote source code and/or documentation for the project. KC, MWF, AG, YVB wrote code to facilitate integration with EMPeror. LP, HCK, SS, ADS, YVB, RK managed the project. KC, MWF, GR, NH, KLB, AT, JMG, LM, APC, NLM, CM, PCD, KGA, LP, YVB analyzed and interpreted the datasets presented in this paper. KC, MWF, GR, DM, NH, KLB, YVB contributed text to the methods section. All the authors contributed to the final version of the manuscript.

## Competing Interests

We declare none.

## Notes

### Competing Interest Statement

The authors have declared no competing interest.

