## Supplementary figures and images for "EMPress enables tree-guided, interactive, and exploratory analyses of multi-omic datasets"

### Supplemental figure 1

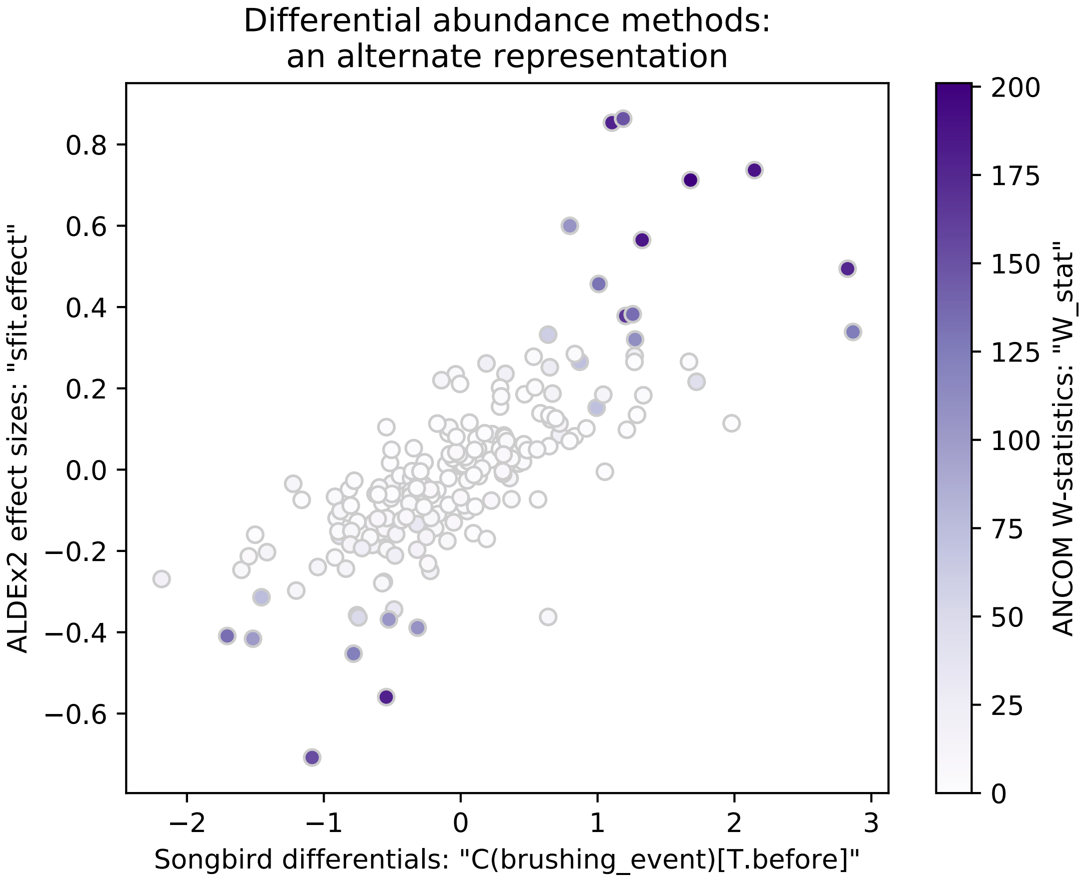
