## Supplemental text for "EMPress enables tree-guided, interactive, and exploratory analyses of multi-omic datasets"

### Methods

All EMPress plots in this paper were visualized and stylized in Safari (14.0) or Google Chrome (85.0.4183.121) using a MacBook Pro (15-inch, 2017) with a 2.9 GHz Quad-Core Intel Core i7 7820QM, 16GB of RAM, a Radeon Pro 560 4GB and Intel HD Graphics 630 1536 MB graphics processor. Analyses, and steps to reproduce the figures in this paper can be found here: https://github.com/knightlab-analyses/empress-analyses

### Earth Microbiome Project

The EMP release 1 [1] table, tree, and metadata were used to generate the visualization (ftp://ftp.microbio.me/emp/release1). The original feature table was subset to remove sterile water blanks and mock community samples. The table contains the taxonomic assignments used to annotate tips. For ease of visualization, only the top 5 most abundant phyla annotations in the dataset were kept while microbial features annotated with any other phylum (or unspecified) were categorized as "Other." A distance matrix was generated by computing the unweighted UniFrac distances between samples [4, 21] that was then used to generate the principal coordinates plot.

A subset of the feature table was generated by extracting all samples in the middle 90% of pH range (4.7 - 9) to remove outliers. Samples without a valid pH value were removed. For each remaining feature, the number of samples (log10) in which this feature occurred and the mean pH of those samples were calculated and saved as a feature metadata file passed into EMPress. Calculations were performed using NumPy v1.18.1 [22] and Pandas v0.25.3 [23, 24]. Distributions of pH were plotted using matplotlib v3.1.3 [25] and seaborn v0.10.0 [26].

### COVID-19 Metatranscriptome Dataset

The COVID-19 bronchoalveolar lavage fluid metatranscriptome sequencing data [17] consists of COVID-19 (n=8), community-acquired pneumonia (n=25), and healthy control samples (n=20). The tree for this dataset corresponds to the KEGG enzyme code (EC) [19] hierarchy. Sequencing reads were processed and annotated with EC feature labels using PRROMenade [27, 29] with a database of bacterial and viral protein domains from the IBM Functional Genomics Platform [28], as previously reported [29]. Differential abundance per feature was determined by performing a K-S test on average RoDEO-processed [29, 30] values per sample. A cutoff ($p < 0.05$) was applied to focus the visualization on the features that are significantly more abundant or less abundant in COVID-19 patients compared to healthy controls and/or community-acquired pneumonia samples.

### Global Food-omics Dataset

The untargeted metabolomics dataset was generated using a QTOF mass spectrometer in positive ionization mode (Bruker). The samples presented in this dataset were processed using Qemistree version 2020.1.1+14.g1b4edb4 running in QIIME2 version 2019.7. For ease of interpretation, the dataset was subset to keep features with a superclass assignment, and keep samples with a *common meal type* classification. The tip barplots show the proportion of samples where each small molecule is present summarized by *meal type*.

### Differential Abundance Comparison of Oral Microbiomes

The oral microbiome 16S rRNA sequencing data used in [18] was re-visualized for Fig. 2(C). This dataset comprises n=32 samples total, taken before and after subjects brushed their teeth.

Some paired samples were taken more than once from the same subject; in total, these 32 samples were contributed by 10 unique subjects.

The sequences in this dataset were processed (in July 2018) using Deblur v1.0.4 [31] through q2-deblur in QIIME 2 2018.6 [2] Taxonomic classifications were assigned (in August 2018) using q2-feature-classifier [32]'s classify-sklearn method [33], using the Greengenes reference database [34], also in QIIME 2 2018.6. To construct a rooted tree from the sequences in this dataset (in September 2020) we used QIIME 2 2019.10's `qiime phylogeny align-to-tree-mafft-fasttree` pipeline [35, 36, 37].

In [18], three differential abundance tools were run on this dataset, using the "brushing_event" metadata field (indicating before/after toothbrushing status) as the sole field across which to identify differentially abundant features. The three differential abundance tools used in [18] and visualized in Fig. 2(C) are Songbird [18], ALDEx2 [38], and ANCOM [39]. Songbird's column of feature differentials (describing the estimated log-fold changes of each feature between the "after" and "before" brushing states) is shown as the innermost barplot layer in Fig. 2(C); ALDEx2's per-feature effect size is shown as the middle layer; and ANCOM's per-feature W-statistic is shown as the outermost layer. For both Songbird and ALDEx2 results, higher values indicate association with before-brushing samples (i.e. features that decreased most from toothbrushing, for example secondary metabolizers present on the outer layers of dental plaque biofilms such as Haemophilus) while lower values indicate association with after-brushing samples (i.e. features that decreased least from toothbrushing, for example primary metabolizers such as Actinomyces that are rooted at the base of the biofilm) [18]. ANCOM's W-statistic corresponds to the number of log-ratio hypothesis tests in which a given feature was found to be differentially abundant between before- and after-brushing samples [39] (https://forum.qiime2.org/t/1844/10). Since Songbird and ALDEx2's results include directionality between before and after brushing, they are shown in Fig. 2(C) with a "diverging" color map; ANCOM's W-statistic does not include this information, and is therefore shown with a "sequential" color map (Supplemental Fig. 1).

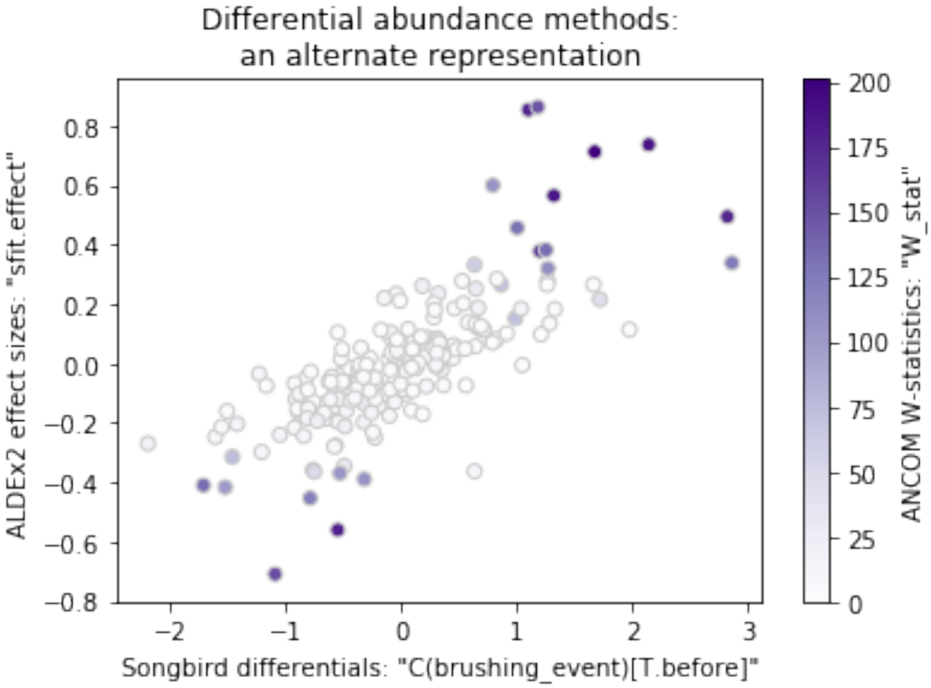


**Supplemental Figure 1. Scatterplot comparing the three differential abundance methods' results shown in Fig. 2(c).** The x-axis and y-axis represent the Songbird differentials and ALDEx2 effect sizes for the features in the dataset, while each feature point is colored by its ANCOM W-statistic. This demonstrates that our starting interpretation of how these results relate—i.e. that Songbird and ALDEx2's results have similar "directionality" between the before- and after- toothbrushing states, and that ANCOM's W-statistic lacks this same directionality—is reasonable. The nine features for which no Songbird differentials were computed are omitted from this plot. This scatterplot was produced using matplotlib [40].

We note that Songbird did not produce differentials for nine features in the dataset; these features were filtered out for reasons such as presence in a relatively small number of samples. These features have no bars drawn for them within the innermost barplot layer, and the color legend shown within Empress contains a warning to this effect.

### Animated Analysis of SARS-CoV-2

The GISAID [41] SARS-CoV-2 genome alignment and genome metadata were obtained on September 21, 2020. Sequences were converted to DNA, and subset to the set of sequences associated with Italy, Madrid, King County, San Diego, Brooklyn, Queens, and Manhattan. Highly gapped and high entropy positions in the alignment were filtered using q2-alignment (2020.6; default parameters). A tree was estimated using FastTree [36] (v2.1.10 compiled with double precision support; default options except -fastest), and subsequently rooted using midpoint rooting as implemented by q2-phylogeny (2020.6; default parameters).

Separately, a sliding window procedure was developed to assess the observed SARS-CoV-2 genomes within a given time period within a geographic location. To do so, the metadata was partitioned into the respective locations (note: the three New York boroughs were treated as New York) and ordered by the genome date information. A sliding window width of seven days was used, and a sample was retained only if five or more strains were observed within a window. These windows were then aggregated into a BIOM table [42] with the GISAID strain identifier on one axis, and a "community sample" identifier on the other. Unweighted UniFrac (q2-diversity 2020.6; [2, 21]) was then computed over these samples followed by a principal coordinates analysis.

### Implementation Details

EMPress is implemented as a QIIME2 plugin capable of generating HTML documents with a self-contained visualization user interface. The code-base is composed of a Python component and a JavaScript component. The Python code-base is responsible for data validation, pre-processing, filtering, and formatting. User interaction, rendering, and figure generation are all handled by the JavaScript code-base. In both cases, we rely on the balanced parentheses data structure [10] to rapidly operate on the tree structures.

EMPress' Python code-base currently uses NumPy [22], SciPy [43], Pandas [23, 24], Click (https://palletsprojects.com/p/click/), Jinja2 (https://jinja.palletsprojects.com/), scikit-bio (http://scikit-bio.org), the BIOM format [42] iow (https://github.com/wasade/improved-octo-waddle) [10 ], and EMPeror [3]. The JavaScript code-base uses Chroma.js (https://gka.github.io/chroma.js/), FileSaver.js (https://github.com/eligrey/FileSaver.js/), glMatrix (http://glmatrix.net/), jQuery (https://jquery.com/), Require.js (https://requirejs.org/), Spectrum (https://bgrins.github.io/spectrum/), and Underscore.js (https://underscorejs.org/). For testing and linting, EMPress' Python code-base uses flake8 (https://flake8.pycqa.org/en/latest/) and nose (https://nose.readthedocs.io/), and EMPress' JavaScript code-base uses QUnit (https://qunitjs.com/), qunit-puppeteer (https://github.com/davidtaylorhq/qunit-puppeteer/), jshint (https://jshint.com/about/), and Prettier (https://prettier.io/).

As of writing, EMPress supports drawing trees using three standard layout algorithms ("rectangular", "circular", and "unrooted"), coloring the tree using sample and feature metadata, collapsing clades based on common metadata values, adding tip-aligned barplots for sample and feature metadata, and summarizing the feature containment within sample groups using interactive node selections.

EMPress' "unrooted" layout algorithm is translated from code from Gneiss [7], which was in turn adapted from PyCogent [44], and is an implementation of the equal-angle algorithm described in [45]. EMPress' "rectangular" and "circular" layout algorithms are adapted from code from TopiaryExplorer [46], and resemble the rooted tree drawing algorithms described in [45]. EMPress also includes the ability to reorder sibling clades in the rectangular and circular layouts by the number of tips contained within each clade; this functionality was inspired by iTOL [15]'s "leaf sorting" option, and uses tree traversal code adapted from scikit-bio (http://scikit-bio.org).

In order to integrate EMPress and EMPeror, we link together events triggered by each of the applications by inserting “callback” code that can be executed in one application when a given event occurs. These events notify each tool that a particular action needs to take place, and if needed what data should be used in this context. For example, when a user selects a group of samples in EMPeror, the “select” event is triggered with a collection of sample objects. EMPress responds to this event by searching for the tips in the tree corresponding to features contained within these samples, and updates the color according to the object’s attributes. The subscription mechanism also enables users to select a node in EMPress to highlight the samples containing this node or one of its descendants in EMPeror, link biplot [47] arrows in EMPeror to nodes in the tree, highlight groups by double-clicking a category in EMPeror’s color legend, and synchronize animated ordinations [48] by coloring the tree according to the current frame on screen.

### Methods only references


1. McDonald, D. *et al.* Striped UniFrac: enabling microbiome analysis at unprecedented scale. *Nat. Methods* **15**, 847–848 (2018).
2. Harris, C. R. *et al.* Array programming with NumPy. *Nature* **585**, 357–362 (2020).
3. McKinney, W. Data Structures for Statistical Computing in Python. in 56–61 (2010). doi:10.25080/Majora-92bf1922-00a.
4. Jeff Reback *et al. pandas-dev/pandas: Pandas 1.1.2*. (Zenodo, 2020). doi:10.5281/zenodo.4019559.
5. Thomas A Caswell *et al. matplotlib/matplotlib v3.1.3*. (Zenodo, 2020). doi:10.5281/zenodo.3633844.
6. Michael Waskom *et al. mwaskom/seaborn: v0.10.0 (January 2020)*. (Zenodo, 2020). doi:10.5281/zenodo.3629446.
7. Utro, F. *et al.* Hierarchically Labeled Database Indexing Allows Scalable Characterization of Microbiomes. *iScience* **23**, 100988 (2020).
8. Seabolt, E. *et al.* IBM Functional Genomics Platform, A Cloud-Based Platform for Studying Microbial Life at Scale. *IEEE/ACM Transactions on Computational Biology and Bioinformatics* 1–1 (2020) doi:10.1109/TCBB.2020.3021231.
9. Haiminen, N., Utro, F., Seabolt, E. & Parida, L. Functional profiling of COVID-19 respiratory tract microbiomes. *bioRxiv* 2020.05.01.073171 (2020) doi:10.1101/2020.05.01.073171.
10. Haiminen, N. *et al.* Comparative exomics of Phalariscultivars under salt stress. *BMC Genomics* **15**, S18 (2014).
11. Amir, A. *et al.* Deblur Rapidly Resolves Single-Nucleotide Community Sequence Patterns. *mSystems* **2**, (2017).
12. Bokulich, N. A. *et al.* Optimizing taxonomic classification of marker-gene amplicon sequences with QIIME 2’s q2-feature-classifier plugin. *Microbiome* **6**, 90 (2018).
13. Pedregosa, F. *et al.* Scikit-learn: Machine Learning in Python. *MACHINE LEARNING IN PYTHON* 6.
14. McDonald, D. *et al.* An improved Greengenes taxonomy with explicit ranks for ecological and evolutionary analyses of bacteria and archaea. *ISME J* **6**, 610–618 (2012).
15. Katoh, K. & Standley, D. M. MAFFT Multiple Sequence Alignment Software Version 7: Improvements in Performance and Usability. *Mol Biol Evol* **30**, 772–780 (2013).
